# The evolution of multicellular complexity: the role of relatedness and environmental constraints

**DOI:** 10.1101/836940

**Authors:** RM Fisher, JZ Shik, JJ Boomsma

**Affiliations:** Section for Ecology and Evolution, Department of Biology, University of Copenhagen, Denmark; Smithsonian Tropical Research Institute, Apartado 0843-03092, Balboa, Ancon, Republic of Panama

**Keywords:** major evolutionary transitions, multicellularity, scaling

## Abstract

A major challenge in evolutionary biology has been to explain the variation in multicellularity across the many independently evolved multicellular lineages, from slime moulds to humans. Social evolution theory has highlighted the key role of relatedness in determining multicellular complexity and obligateness, however there is a need to extend this to a broader perspective incorporating the role of the environment. In this paper, we formally test Bonner’s 1998 hypothesis that the environment is crucial in determining the course of multicellular evolution, with aggregative multicellularity evolving more frequently on land and clonal multicellularity more frequently in water. Using a combination of scaling theory and phylogenetic comparative analyses, we describe multicellular organisational complexity across 139 species spanning 14 independent transitions to multicellularity and investigate the role of the environment in determining multicellular group formation and in imposing constraints on multicellular evolution. Our results, showing that the physical environment has impacted the way in which multicellular groups form, could shed light on the role of the environment for other major evolutionary transitions.

## Introduction

Macroscopic life on earth has been shaped by the evolution of multicellularity from unicellular ancestors. Multicellularity is a complex and variable trait, ranging from simple cell aggregations found in yeast to differentiated metazoan organisms, with much diversity in between [1]. For example, our bodies contain 10^14^ cells with more than 200 specialized types [2] but *Volvox* is 10 orders of magnitude smaller and has just 2 cell types [3]. Some lineages have become obligately multicellular, where cells only exist as part of a multicellular organism (e.g. animals), whereas others remain facultative, switching between a unicellular and multicellular lifestyle (e.g. cellular slime moulds) [4].

A major challenge in evolutionary biology has been to explain this variation in complexity among multicellular lineages. Social evolution theory has greatly advanced our understanding of the evolution of multicellularity, primarily through clarifying the factors that favour the cooperation needed to become multicellular. We understand how relatedness between cells is crucial in determining when altruism can evolve (for example, in *Dictyostelium* slime moulds) [5], division of labour between cell types [6] and the proliferation of cheaters [7–9]. It has also become clear that clonal relatedness (r = 1) is a necessary, albeit not sufficient, condition for the evolution of obligate multicellularity like we see in animals and plants, and that these lineages have more cell types than those with facultative multicellularity [4].

However, there are limits to the variation in multicellular complexity that is explained by relatedness. For example, both land plants and fungi have cells that are clonal and obligately multicellular, but plants have approximately 10 times more cell types than fungi [10] and it is unclear what can explain these differences. There are good reasons to speculate that the environment could be an important factor shaping the first trajectories of multicellular evolution with lasting consequences for later elaborations. Firstly, the environment itself could determine the way in which multicellular groups form and hence relatedness between cells. Bonner (1998) observed that clonal group formation, where daughter cells remain attached to mother cells after division, seems to be more common in lineages that originated in the sea compared to species that originated on land [11]. If this is the case, it would mean that the environment where multicellularity originates could have profound consequences for subsequent evolutionary possibilities. Secondly, the physical constraints associated with living in water or on land are likely to affect many aspects of phenotypic evolution, for example the need for support and structural reinforcement tissues, the diversity of dispersal mechanisms, and the sustaining the biomechanics of active motility.

Scaling theories provide powerful tools to test for such constraints, since an organism’s body size can accurately predict complex traits such as metabolic rate, lifespan, and growth rate [12], and since the shapes of these relationships reflect fundamental physiological constraints on how diverse organisms can evolve [13]. Scaling relationships can also reveal outlier taxa that highlight cases where evolutionary innovation fueled the breaking of ecological and physiological constraints [14]. In practice, scaling parameters (*i.e*. the slope *(b)* and intercept (*a*) in the equation y = *aM^b^*) represent mechanistic hypotheses that, for the purpose of this study, relate the number of cell types (y) to the total number of cells (*M*). Isometric scaling (*b* = 1) provides a null model, predicting that cell type and cell number increase at the same rate (every added cell is a new type), and allometric scaling (*b* < 1) would indicate that cell type increases at a slower rate than cell number, such that small organisms have more cell types relative to their body size.

There is a need to build on our understanding of the fundamental factors influencing multicellular evolution – primarily the role of relatedness – and extend this to a broader perspective incorporating the role of the environment. The objectives of this paper are to: (1) describe the variation in multicellular organisational complexity across 139 species by investigating the scaling relationships between body size (total number of cells) and number of cell types; (2) use phylogenetically-controlled comparative analyses across 14 independent multicellular transitions to assess the extent to which the environment determines how multicellular groups form and the consequences for whether obligate multicellularity evolves; and (3) test whether constraints imposed by the environment can explain why some lineages have reached higher levels of organisational complexity than others, and can account for part of the variation in cell-type diversity and differences in scaling relationships. We use the term organisational complexity to highlight that division of labour is fundamental to multicellularity, that the number of cell types is a marker of division of labour and that any form of division of labour requires organisational integration.

## Results

### Describing variation in body size and complexity

Across 139 species, representing 14 independent transitions to multicellularity, the scaling of cell type and cell number is strongly allometric (reduced major axis regression, RMA): slope = 0.14, CI = 0.13 - 0.16, R^2^ = 0.64, Figure 1a & b). This means that despite a positive association, the number of cell types increases much more slowly with cell number than arithmetic proportionality (i.e. isometric scaling) would predict. In other words, small organisms are organisationally more complex for their size than large organisms.

**Figure 1.**
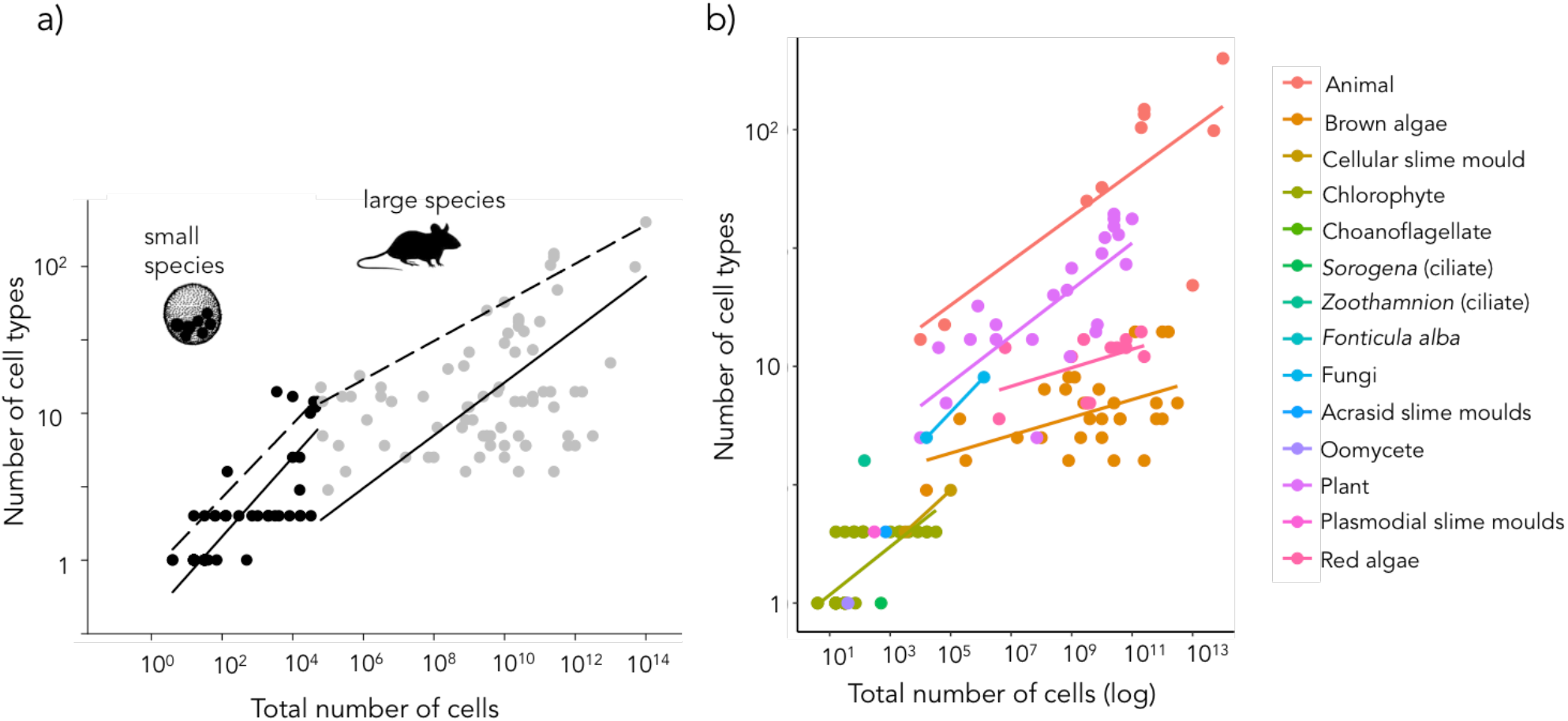
**(a) Scaling across multicellular organisms.** The relationship between number of cell types and total number of cells for small (in black, between 4 – 10^4^ cells) and large multicellular species (in grey, between 10^4^ – 10^14^ cells) shown on logarithmic axes. Small species show a steeper allometry (reduced major axis regression: slope = 0.27 (CI 0.23 −0.33) compared to large species (reduced major axis regression: slope = 0.18 (CI 0.15 – 0.23). Solid lines show the reduced major axis regression and dashed lines show regressions through the upper 90% quantile of the data. We estimated the breakpoint of 4.8 ±0.9 (corresponding to 6.3 x 10^4^ total cells) using the ‘segmented’ package in R. **(b) Multicellular organisational complexity across different multicellular lineages.** Organisational complexity, measured as both the number of cell types and the total number of cells, for each of the independently evolved multicellular lineages. These data have been taken from the dataset of Bell & Mooers (1997) [10].Original data are from the data set of Bell & Mooers (1997) and images of *Mus musculus* and *Volvox* are from Phylopic (http://phylopic.org/). The statistical results of the different regressions are given in Table S1.

We next found that there was a difference in the scaling relationship between number of cell types and total number of cells for small versus large species (Table S1, S2). Specifically, we identified the estimated breakpoint in the regression as 6.3 x 10^4^ total number of cells, corresponding to 4.8 ± 0.9 cell types, where the scaling relationship changes. Small species (before the breakpoint) showed an allometric slope about twice as steep (RMA: slope = 0.27, CI = 0.23 – 0.33, R^2^ = 0.61) as large species (above the breakpoint) (RMA: slope = 0.18, CI = 0.15 – 0.23, R^2^ = 0.15). This implies that larger organisms face a unique set of more stringent constraints on the accumulation of new cell types than small organisms, and supports the observation that lineages consisting of small species gain new cell types more quickly as they grow in size.

We further sought to understand the substantial reduction in cell type variation explained by cell number for large organisms (i.e. R^2^ = 0.15) using a technique called quantile regression. Regression through the upper 90% quantile of the dataset suggests that there is an upper threshold to the number of cell types a species can have for its size, whereas there is a lot of variation in the number of cell types below that threshold (Figure 1a, dashed lines). This suggests that: there could be other factors limiting the number of cell types below that threshold and these other limiting factors are especially important in larger species since the slope describing this upper limit is far shallower (*b* = 0.13) than the upper limit for small species (*b* = 0.25) (Table S2).

### The origins of multicellularity in different environments

Our results show that the physical environment (whether or not a species lives in the water or on land) has had a major impact on both the origins and subsequent elaborations of multicellularity, both in determining how multicellular groups originally form and how organisational complexity subsequently evolves.

We found that lineages in aquatic environments were significantly more likely to form multicellular groups through daughter cells remaining attached to mother cells after division (clonal group formation) (MCMCglmm, difference between aquatic & terrestrial: posterior mode = 5.74, credible intervals (CI) = 2.91 – 9.79, p_diff_ = 0.0008, N_species_ = 139, Figure 2a). All of the multicellular lineages in our dataset that have their origins in water form multicellular groups in this way, whereas two thirds of the lineages that originated on land form groups through aggregation (non-clonal group formation) (Figure 2a). In fact, there are only two lineages that originated on land that employ clonal group formation – the Fungi and the plasmodial slime moulds and these tend to grow in terrestrial environments of saturated humidity. This result confirms Bonner’s original observation that clonal group formation is more common in multicellular lineages originating in the sea [11].

**Figure 2:**
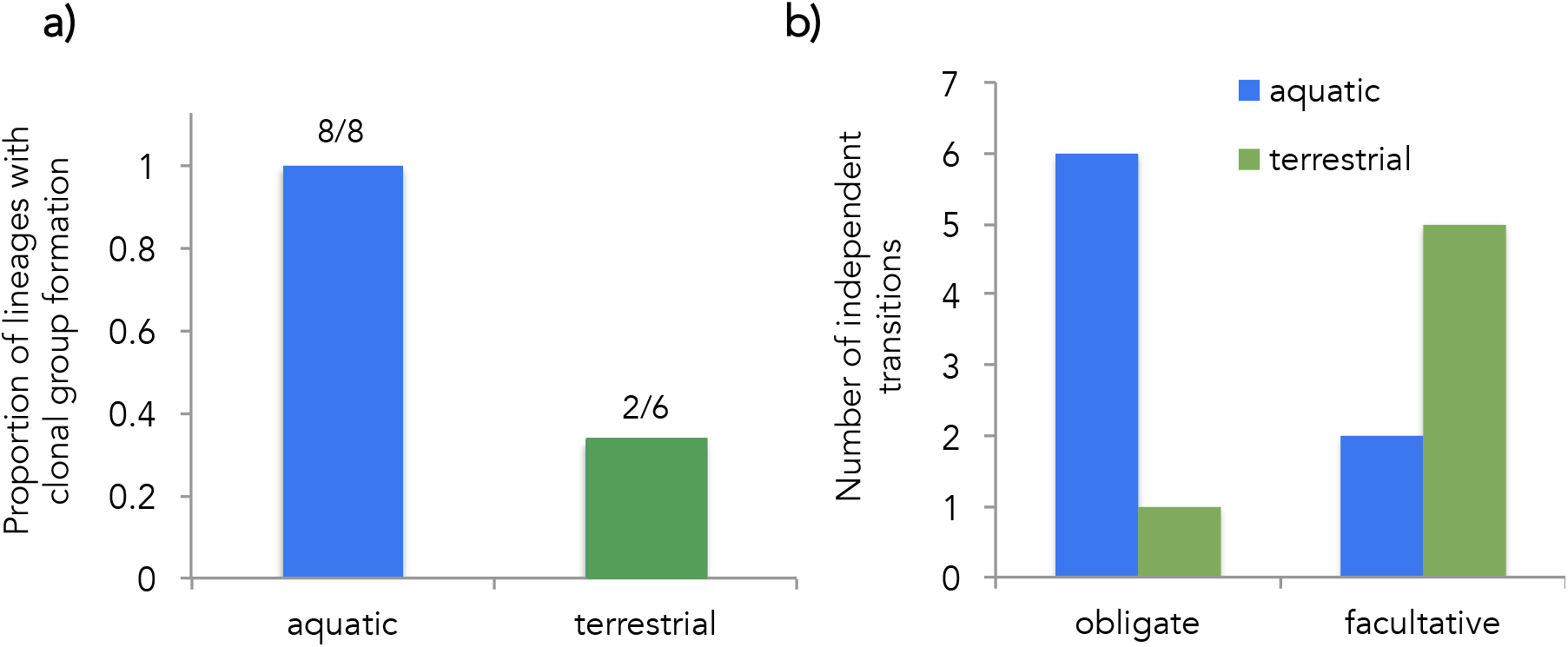
The origins of multicellularity in different environments. **(a)** The proportion of lineages that have clonal group formation that originated in aquatic and terrestrial environments. All multicellular lineages that originated in the sea have clonal group formation (8/8 lineages) whereas most of the multicellular lineages that originated on land have non-clonal group formation (4/6 lineages). **(b)** Multicellular lineages that originated in water more commonly evolve obligate multicellularity (6/8 lineages) compared to lineages that originated on land, which more often remain facultatively multicellular (5/6 lineages).

Secondly, we found that the transition to obligate multicellularity was significantly more likely to occur in aquatic environments compared to on land. Most (5 of 6) lineages that evolved multicellularity on land remained facultatively multicellular (difference between aquatic & terrestrial: posterior mode = 6.59, CI = 4.29 – 8.72, p_diff_ = < 0.0001, N_species_ = 139, Figure 2b). The only multicellular lineage that has evolved obligate multicellularity on land is the Fungi. This is consistent with this lineage also being a rare example of clonal group formation that originated on land, as the resulting clonal relatedness between cells is significantly associated with the transition to obligate multicellularity [4].

### Multicellular organisational complexity on land versus in water

We found that the number of cell types of multicellular species currently found on land was significantly higher than those currently found in aquatic environments (Figure 3a), whilst controlling for the total number of cells (posterior mode = −0.77, CI = −1.42 to −0.11, p_diff_ = 0.02, N_species_ = 137, Figure 3b). The average number of cell types for aquatic lineages is 8 whereas for terrestrial lineages it is 25. Species on land were however not significantly larger in size than those found in the sea (posterior mode = −2.79, CI = −9.04 to 1.81, p_diff_ = 0.12, N_species_ = 137). Overall, there was a significant phylogenetic correlation between number of cell types and total number of cells, meaning that species with more cell types also tend to be bigger due to their shared ancestry (posterior mode = 0.90, CI = 0.72 to 0.96, p_diff_ = < 0.0001, N_species_ = 137). However, we also found a significant phenotypic correlation between these two variables, meaning that the association is also a result of a shared environment (posterior mode = 0.56, CI = 0.19 to 0.76, p_diff_ = 0.004, N_species_ = 137).

**Figure 3:**
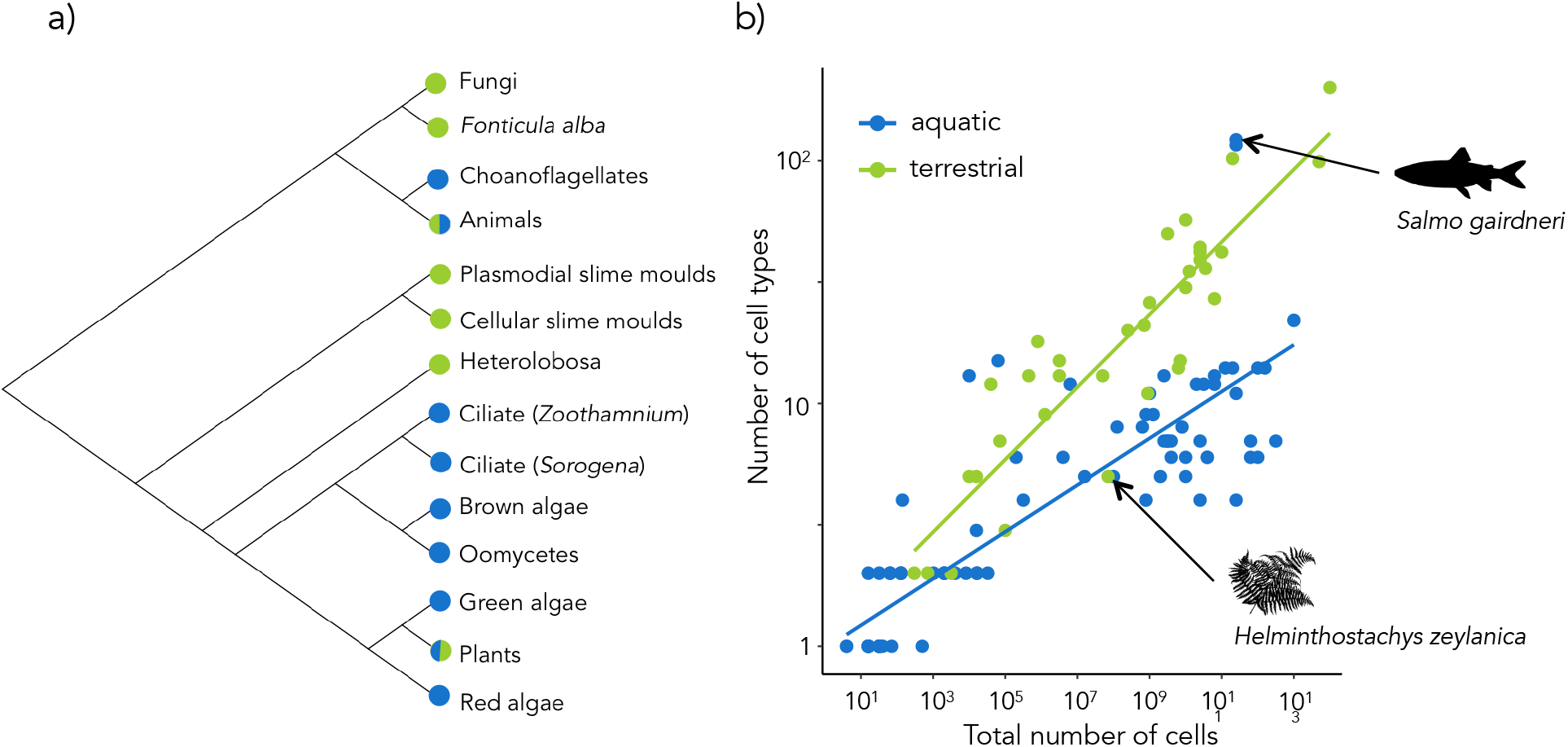
Organisational complexity and environmental constraints. **a)** A summarised phylogram of the lineages that have independently evolved multicellularity. The current environment is shown as terrestrial (in green), aquatic (in blue) or both (half green, half blue) for species that have a substantial number of species in both environments. **b)** Multicellular complexity, measured as both the number of cell types and the total number of cells, for species currently found in aquatic and terrestrial environments. Two notable outliers (*Salmo gairdneri* and *Helminthostachys zeylanica*) are highlighted with black arrows and images and further interpreted in the Discussion. These data have been taken from the dataset of [10].

## Discussion

We were interested in how multicellular complexity scales with body size and the role the physical environment could play in shaping the course of multicellular evolution. Overall, we found that the number of cell types scales allometrically with the total number of cells (echoing Bonner’s observations [1]), and that the specific scaling relationship is different for small versus large species. Our comparative analyses also show that the environment (aquatic or terrestrial) has a crucial impact on the trajectory of multicellular evolution. Firstly, we found that clonal group formation giving rise to obligate multicellularity has been significantly more common in lineages that evolved in aquatic environments. Secondly, we showed that current environmental conditions have an impact on multicellular evolution, with species living on land having a higher number of cell types compared to species found in aquatic environments.

Bonner (1998) observed that clonal group formation was more common in multicellular lineages that evolved in the sea whereas aggregation was more common in terrestrial lineages. Our results provide formal support for this observation by including additional lineages and using phylogenetically controlled comparative analyses. Bonner speculated that this pattern could be because of water currents, meaning that cells in water need to stick together after they divide if they want to reap the benefits of being in a group [11]. This is not the case on land, where cells must use active motility (e.g. cilia, flagella, amoeboid movement) in order to form multicellular groups. It is clear from the species in our dataset that the multicellular lineages found on land (*Dictyostelium, Sorogena, Physarum, Fonticula*) all have some form of motile cell stage, and even the Fungi have an ancestral lineage with motile cells [31]. Therefore, it seems plausible that the biophysics of moving through air and water has had a profound impact on the way in which multicellular groups could form on land and in the sea.

It is likely that our dataset underestimates the number of lineages that have facultative multicellularity. These species have transient multicellular phenotypes and so many species are still identified as unicellular. For example, *Saccharomyces cerevisiae* has a variety of multicellular phenotypes [32] and yet is still often not recognised as being facultatively multicellular [33]. Other lineages, notably the green algae, have evolved facultative multicellularity many times [21] and there are likely new examples to be found on land as well. However, it is unlikely that we have underestimated the number of lineages with obligate multicellularity. This is because these species tend to be bigger and therefore more visible and complex [4] and potentially better studied [18,34]. There is no obvious reason to assume that under- or overestimation would be biased towards terrestrial or aquatic species. By inflating the number of facultative lineages, we would therefore not alter the pattern and the result we find – that obligate multicellularity has evolved much more often in water compared to on land.

Our study reveals a number of intriguing outliers. Firstly, not all species fit the overall pattern of higher organizational complexity on land. There are outlier species in our dataset, including several living in aquatic environments that display levels of complexity more similar to terrestrial species. For example, *Salmo gairdneri* (Rainbow trout) has an estimated 116 cell types and a total size of 2.51 x 10^11^ cells, which is more similar to the terrestrial *Mus musculus* (mouse) than to other aquatic species. Another example is *Helminthostachys zeylandica* (a member of the fern family) that has much lower complexity for its size than other terrestrial multicellular species (just 5 cell types) (Figure 3b). Secondly, a major and strikingly unusual lineage are the Fungi, that develop multicellularity through clonal group formation but display (mostly) simple multicellularity. However, our dataset only included 3 species from Kingdom Fungi and there is also a lack of data on fungal multicellularity in the wider literature. Perhaps a closer look at the Fungi as ‘exceptions to the rule’ could help to unravel the relationship between the environment and multicellular complexity.

Not only does the environment affect how multicellular groups form, but we show that it also has a major impact on the scaling relationships between size and complexity. Species that live on land tend to be more complex for their size compared to species that live in water (i.e., with a higher slope, Figure 3b) and this could be for several reasons. Land dwelling organisms need more support structures than their aquatic counterparts – this is because water provides natural support through buoyancy whereas air does not. Organisms living on land therefore needed to increasingly invest in stems and skeletons to ‘hold themselves’ up as their body size increases (i.e. skeleton mass ~ *M^b > 1^*, [12]), possible leading to greater diversification of cell types and tissues than organisms in the sea. This scaling logic can further be extended to resource allocation dynamics within organisms (e.g., vascular networks,), although systematic effects on cell diversity and differences between land and water remain to be elucidated. There are also other potentially confounding physiological parameters, for example that autotrophic lineages compete for light, both on land and in the water, whereas heterotrophic lineages do not, but that support tissues on land are more costly to maintain (e.g. rain forest trees versus kelp forest).

The parallel evolutionary events towards obligate multicellularity are examples of major evolutionary transitions in individuality [35]. A key aim of major transition research is to identify common patterns across different transitions (e.g. the evolution of prokaryote and eukaryote cells, obligately multicellular organisms, and colonial superorganisms). The fact that these all arose through clonal group formation or as full sibling families (initiated by strictly monogamous pairs) implied that reproductive allocation conflicts did not play a role, as they usually do in promiscuous or chimeric associations that do not make such major transitions [36,37] [4,38]. Strict vertical transmission of symbionts, including mitochondria and plastids, was also a potent force to avoid conflict [39]. Our results, showing that the physical environment has impacted the way in which multicellular groups form, could therefore shed light on the role of the environment for other major evolutionary transitions. For example, how have physical conditions across nesting habitats (e.g. subterranean versus arboreal; [40]) influenced the necessary and sufficient conditions for insect colonies to commit to obligate division of labour via specialized and physically differentiated castes?

## Material and Methods

### Data collection

The data used in this study were originally published in Fisher *et al*. (2013) and are stored in the data depository Dryad (original data can be found here: https://datadryad.org/resource/doi:10.5061/dryad.27q59). In summary, we conducted an extensive literature search on multicellular species, searching specifically for information on multicellular complexity, the ways in which groups formed and whether or not they were obligately or facultatively multicellular. We used data from Bell & Mooers (1997) on the number of cell types and total number of cells to estimate multicellular complexity for each multicellular species [10] as this is the most taxonomically-representative dataset on cell types, to our knowledge [41]. Our full dataset can be found in Table S6.

In this study, we expanded this original dataset by adding information on the ancestral and current environment of each species. We considered any species found on land as terrestrial and any species found in freshwater, brackish or marine environments as aquatic. We found information about the current environment of a species by searching on Google Scholar for publications and also taxa-specific websites, such as AlgaeBase and WoRMs. Where there was only information about ancestral or current environment at a higher taxonomic level (i.e. at the family level but no generic or species information) we assumed it was the same environment for the species in our dataset. We found information on the ancestral environment of each species through broad reviews on the origins of multicellularity including Bonner 1998, Knoll 2011 & Umen 2014 [11,21,28]. It is important to stress that we were interested in the ancestral environment *when multicellularity evolved* and therefore that was not always the same as the ancestral environment for the whole lineage, including unicellular groups (e.g. for the Fungi, James *et al*. 2006).

Of the 139 species in the dataset, 18 species had a terrestrial ancestral environment and 121 species had an aquatic ancestral environment. For the current environment, 84 species are aquatic, 43 are terrestrial and 12 are unknown.

### Independent transitions to multicellularity

Using information from published papers, we identified that within the eukaryotes there have been at least 14 independent transitions to multicellularity (both facultative and obligate) (Table 1, Figure 3a). However, we have most likely underestimated the number of transitions in several groups due to uncertainty about the number of independent transitions within them. For example, it is thought that there have been at least 2 transitions to obligate multicellularity within the Fungi [27,28] and many transitions to facultative multicellularity in the green algae [21] and in the red algae [18]. Therefore, our analyses are conservative and assumed just 1 transition within each group.

**Table 1:**
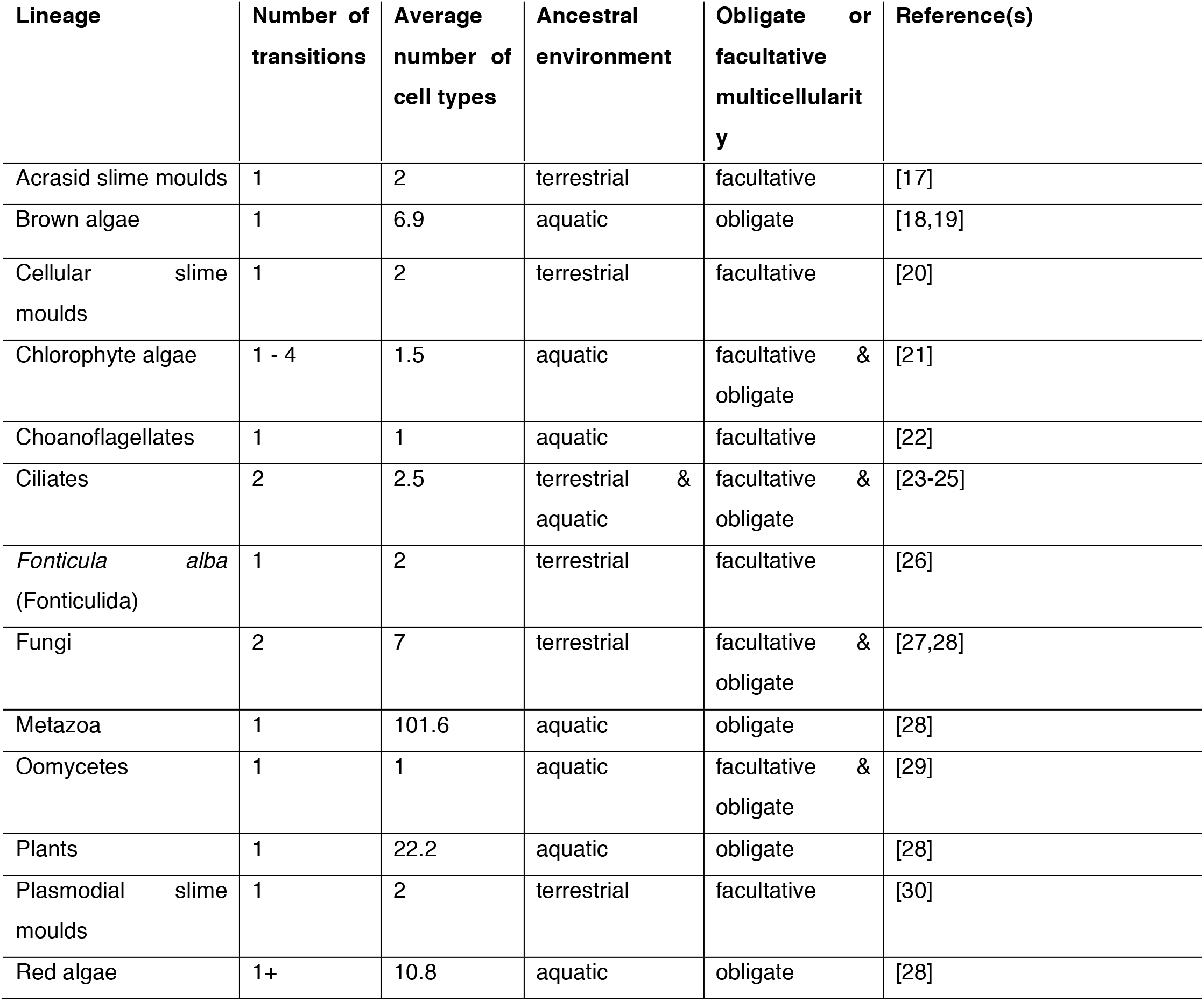
At least 14 transitions to multicellularity occurred within the eukaryotes. Estimates of the number of independent transitions in each lineage are given along with the environment where the lineage originated, average number of cell types, and the corresponding references. We have not included two other known transitions to multicellularity – the diatoms [15] and *Sorodiplophrys* (Stramenopiles) [16] – due to a lack of data on cell types and environment of origin. See Figure 3a.

## Statistical Methods

### Scaling relationships

As a first step in analyzing the data we began with a least square regression to estimate *a* and *b* in the scaling equation log_10_*y* = log_10_*a* + *b*log_10_*M* and describe nature of the dependence of the number of cell types on the total number of cells. We used the R package ‘lmodel2’, we used reduced major axis (RMA) regression to estimate the intercept and slope in the scaling of log_10_(cell type) against log_10_(cell number) across all data, for small species and for large species. RMA is an appropriate line-fitting method in cases when measurement of both Y and X variables are potentially associated with systematic error (e.g. the probability that cell number was precisely measured decreased with increasing body sizes) [42]. RMA (also known as standardized major axis regression) equally weights distances from the regression line in both X and Y directions, with the major axis reflecting the first principal components axis yielded by the covariance matrix, and fitted through the centroid of the data [42]. We then used the package ‘segmented’ in R [43] to test if there is a ‘breakpoint’ in the regression – the point at which the shape of the relationship changes dramatically. This allowed us to estimate the different scaling relationships of small versus larger multicellular species.

We also noted that the scaling relationships appeared triangular and thus hypothesized that they reflect a constraint function such that total number of cells is necessary, but not sufficient to explain variation in number of cell types [44,45]. To test this hypothesis, we used least absolute deviation regression to describe scaling for the upper ninetieth quantiles of the overall plot and separately for the small and large taxa plots [46,47].

### Bayesian analyses

We used the statistical package MCMCglmm [48] to run Bayesian general linear models with Markov Chain Monte Carlo (MCMC) estimation. We fitted three models. Firstly, we tested whether the environment affected the way in which multicellular groups form by fitting a model with group formation as a categorical response variable and the ancestral environment as a categorical explanatory variable (Table S1). Secondly, we tested whether the environment affected the likelihood of obligate or facultative multicellularity by fitting a model with obligate/facultative as a categorical response variable and the ancestral environment as a categorical explanatory variable (Table S3).

Finally, we tested whether multicellular complexity differed between lineages living on the land versus in aquatic environments by fitting a multi-response model with several explanatory variables using the number of cell types and the logarithm of total number of cells as poisson and Gaussian response variables respectively (Table S4). This allowed us to use both number of cell types and the total number of cells as a combined measure of multicellular complexity, rather than having to run several analyses using different response variables. We fitted several categorical fixed effects: the current environment (aquatic or terrestrial), whether the species is obligately or facultatively multicellular, and the mode of group formation (non-clonal or clonal) to control for the known effects of group formation and obligateness on complexity [4].

In the first two models, we used uninformative inverse-gamma priors because we had a categorical response variable. We also fixed the residual variance to 1 and specified family = categorical. In the final model, we used uninformative priors because we had a multi-response model with both poisson and Gaussian response variables and categorical explanatory variables. We ran the models for 6000000 iterations, with a burn-in of 1000000 and a thinning interval of 1000. These were the values that optimised the chain length whilst also allowing our models to converge, which we assessed visually using VCV traceplots. We then ran each model three times and used the Gelman-Rubin diagnostic to quantitatively check for convergence. We showed our models had converged when the PSR was < 1.1.

We calculated the correlations between the number of cell types and the total number of cells (cov(number of cell types, total number of cells)/sqrt(var(number of cell types)* var(total number of cells)) for species in different environments. We tested if the correlation was significantly different between environments by examining if the 95% credible interval of the difference between the correlations spanned 0, and calculating the % of iterations where the correlation for species living in aquatic environments was greater than that for those living on land.

### Phylogeny construction

We built the phylogeny for this study using the Open Tree of Life (opentreeoflife.org), which creates synthetic trees built from published phylogenies and taxonomic information. We then used the R package ‘rotl’ that interacts with the online database and constructs phylogenies (https://cran.r-project.org/web/packages/rotl/index.html). For the majority of species in our dataset, the exact species was also present in a published phylogeny and so we could use phylogenetic information about that species. However, for a few species that were not present in the Open Tree of Life dataset, we had to assign instead a closely related species in the same genera or use a family-level classification. Due to the fact that most species in our dataset represent distant groups on the eukaryotic tree and our phylogeny does not include branch lengths, we were confident this compromise did not affect our statistical analysis.

## Acknowledgements

We thank Stuart West and Guy Cooper for thought-provoking discussions and comments and Stefania Kapsetaki, Jordan Okie, and Jamie Gillooly for helpful edits on a final version of the manuscript. We dedicate this paper to John Tyler Bonner.

