## Supplemental Information for "The evolution of multicellular complexity: the role of relatedness and environmental constraints"

### Supplementary Information and Figures

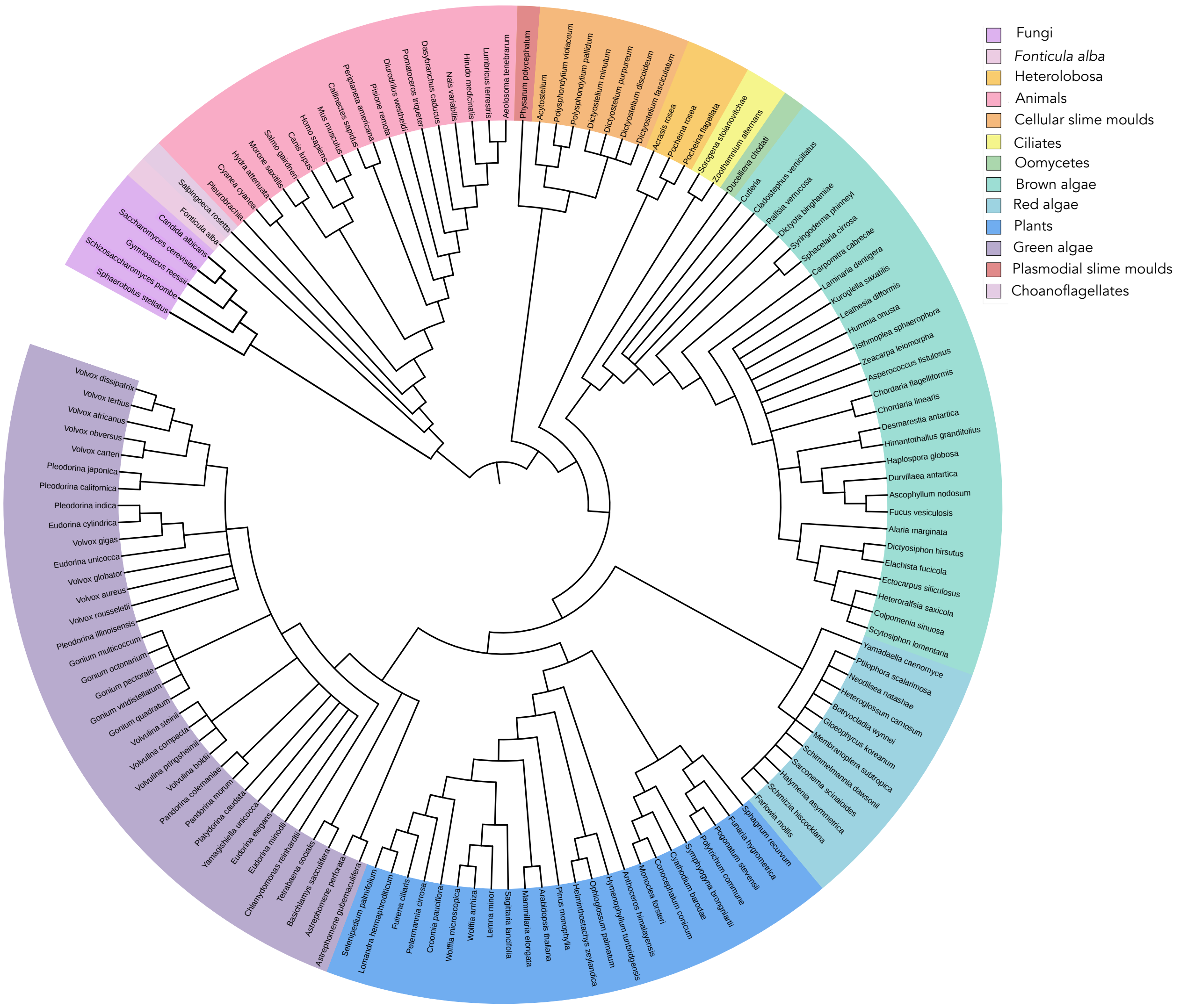


**Figure S1: Phylogeny of the multicellular lineages in our dataset.** Phylogeny created using the Open Tree of Life and 'rotl' package in R and edited using the online software Interactive Tree of Life. Each lineage that independently evolved multicellularity within the Eukaryotes is highlighted in a different colour and the species that appear in our dataset are given at the tips.

### Supplementary Statistical Results Tables

**Table S1:** Statistical results from Least squares (OLS) and Reduced Major Axis (RMA) Regression on the full dataset, small species and large species. *slopes are significantly different from 0

| Model | N | Intercept (2.5% - 97.5% confidence intervals) | Slope (2.5% - 97.5% confidence intervals) | $R^2$ | p value |
|---|---|---|---|---|---|
| OLS (all) | 126 | 0.03 (-0.09 – 0.15) | 0.11 (0.10 – 0.13) | 0.64 | 0.01* |
| OLS (small) | 50 | -0.23 (-0.37 – 0.09) | 0.21 (0.16 – 0.26) | 0.61 | 0.01* |
| OLS (large) | 76 | 0.46 (0.08 – 0.83) | 0.07 (0.03 – 0.11) | 0.15 | 0.01* |
| RMA (all) | 126 | -0.16 (-0.27 - -0.06) | 0.14 (0.13 – 0.16) | *NA* | *NA* |
| RMA (small) | 50 | -0.38 (-0.52 - -0.27) | 0.27 (0.23 – 0.33) | *NA* | *NA* |
| RMA (large) | 76 | -0.59 (-1.00 - -0.26) | 0.18 (0.15 – 0.23) | *NA* | *NA* |

**Table S2 :** Statistical results from the 90% Regression on the small and large species.

| Model | N | Intercept (2.5% - 97.5% confidence intervals) | Slope (2.5% - 97.5% confidence intervals) |
|---|---|---|---|
| 90% Quantile regression (small) | 126 | -0.08 (-0.50 – 0.35) | 0.25 (0.10 – 0.40) |
| 90% Quantile regression (large) | 76 | 0.44 (0.13 – 0.76) | 0.13 (0.10 – 0.17) |

**Table S3:** Analysis of the effect of the ancestral environment on the mode of multicellular group formation, taking into account phylogenetic relationships using MCMCglmm.

| **Response:** Mode of group formation | | | | |
|---|---|---|---|---|
| **Fixed effect** | | **N** | **Posterior mode (CI)** | **pMCMC** |
| Ancestral environment | Aquatic | | 4.80 (1.26 – 7.58) | |
| | Terrestrial | | -1.43 (-4.78 – 0.80) | |
| | Difference | | 5.74 (2.91 – 9.79) | 0.0008 |

**Table S4:** Analysis of the effect of the ancestral environment on whether a species is obligately or facultatively multicellular, taking into account phylogenetic relationships using MCMCglmm.

**Response:** Obligate or facultative

| **Fixed effect** | | **N** | **Posterior mode (CI)** | **pMCMC** |
|---|---|---|---|---|
| Ancestral environment | Aquatic | | 4.04 (2.39 – 5.97) | |
| | Terrestrial | | -2.02 (-4.09 - -0.49) | |
| | Difference | | 6.59 (4.29 – 8.72) | < 0.0001 |

**Table S5:** Analysis of the effect of the current environment, whether the species is obligately or facultatively multicellular and the mode of group formation on the number of cell types and the total number of cells, taking into account phylogenetic relationships using MCMCglmm.

**Response:** The number of cell types

| **Fixed Effect** | | **N** | **Posterior mode (CI)** | **pMCMC** |
|---|---|---|---|---|
| Current environment | Aquatic | | -0.11 (-1.56 – 1.70) | |
| | Terrestrial | | 0.53 (-0.72 – 2.36) | |
| | Difference | | -0.77 (-1.42 - -0.11) | 0.02 |
| Obligate or facultative | Obligate | | 1.97 (-0.36 – 3.87) | |
| | Facultative | | 0.38 (-1.49 – 1.77) | |
| | Difference | | 1.68 (0.06 – 3.24) | 0.02 |
| Mode of group formation | Clonal | | 0.11 (-1.75 – 2.33) | |
| | Non-clonal | | 0.18 (-1.46 – 1.82) | |
| | Difference | | -0.19 (-1.66 – 1.43) | 0.43 |

**Response:** Total number of cells

| **Fixed Effect** | | **N** | **Posterior mode** | **pMCMC** |
|---|---|---|---|---|
| Current environment | Aquatic | | 9.80 (-5.03 – 20.82) | |
| | Terrestrial | | 13.21 (-2.16 – 23.22) | |

| | | | | |
|---|---|---|---|---|
| | Difference | | -2.79 (-9.04 – 1.81) | 0.12 |
| Obligate or facultative | Obligate | | 7.95 (0.94 – 12.97) | |
| | Facultative | | 3.40 (-1.66 – 9.01) | |
| | Difference | | 3.89 (0.65 – 6.62) | 0.007 |
| Mode of group formation | Clonal | | 1.78 (-3.36 – 7.71) | |
| | Non-clonal | | 3.23 (-1.83 – 9.15) | |
| | Difference | | 1.46 (-1.98 – 4.51) | 0.23 |

**Correlations**

| | | | |
|---|---|---|---|
| Number of cell types : Total number of cells | Phylogenetic correlation | 0.90 (0.72 – 0.96) | < 0.0001 |
| | Phenotypic correlation | 0.56 (0.19 – 0.76) | 0.004 |
