## Supplemental data for "The evolution of multicellular complexity: the role of relatedness and environmental constraints"

**Table S6:** The full dataset, with corresponding references, for all of the data used in our analyses.

| Species | Obigate or faculative | Number of cell types | Total number of cells | Group formation | Current environment | Ancestral environment | Reference(s) |
|---|---|---|---|---|---|---|---|
| *Acrasis rosea* | facultative | 2 | 707.9457844 | non-clonal | terrestrial | terrestrial | Raper 1984, Bonner 2003, Brown *et al.* 2011 |
| *Acytostelium* | facultative | 1 | NA | non-clonal | terrestrial | terrestrial | Bonner 2003, Bourke 2011 |
| *Aelosoma tenebrarum* | obligate | 12 | 50118.72 | clonal | NA | aquatic | Brace 1901 |
| *Alaria marginata* | obligate | 14 | 1.00E+12 | clonal | aquatic | aquatic | Kain 1979, Charrier *et al.* 2007 |
| *Anthoceros himalayensis* | obligate | 12 | 39810.71706 | clonal | terrestrial | aquatic | Henra & Handoo 1953, Nishiyama 2007 |
| *Arabidopsis thaliana* | obligate | 30 | | clonal | terrestrial | aquatic | Carroll 2001, Lin *et al.* 1999 |
| *Ascophyllum nodosum* | obligate | 6 | 6.31E+11 | clonal | aquatic | aquatic | Rawlence 1978, Charrier *et al.* 2007 |
| *Asperococcus fistulosus* | obligate | 5 | 10000000000 | clonal | aquatic | aquatic | Bold & Wynne 1978, Charrier *et al.* 2007 |
| *Astrephomene gubernaculifera* | obligate | 2 | 64 | clonal | aquatic | aquatic | Stein 1958, Herron & Michod 2008, Hallmann 2011 |
| *Astrephomene perforata* | obligate | 2 | 64 | clonal | aquatic | aquatic | Herron & Michod 2008, Hallmann 2011 |
| *Basichlamys sacculifera* | obligate | 1 | 4 | clonal | aquatic | aquatic | Nozaki *et al.* 1996, Hallmann 2011 |
| *Beckerlla scalaramosa* | obligate | 12 | 31622776600 | clonal | aquatic | aquatic | Kraft 1976, Graham 1985 |
| *Botryocladia wynnei* | obligate | 6 | 3981071.706 | clonal | aquatic | aquatic | Ballantine 1985, Graham 1985 |
| *Callinectes sapidus* | obligate | 69 | 3.16E+11 | clonal | NA | aquatic | Johnson 1980 |
| *Candida albicans* | facultative | 3 | NA | clonal | NA | terrestrial | Engelberg *et al.* 1998, Whiteway & Bachewich 2007 |
| *Canis familiaris* | obligate | 99 | 5.01E+13 | clonal | terrestrial | aquatic | Adam et al 1983 |
| *Carpomitra cabrecae* | obligate | 7 | 2511886432 | clonal | aquatic | aquatic | Motomura *et al.* 1985, Charrier *et al.* 2007 |
| *Chlamydomonas reinhardtii* | facultative | 1 | 71 | clonal | aquatic | aquatic | Bold & Wynne 1985, Kirk 1998, Lurling *et al.* 2006, Becks *et al.* 2010 |
| *Chordaria flagelliformis* | obligate | 6 | 3981071706 | clonal | aquatic | aquatic | Kornmann 1962, Charrier |

| | | | | | | | |
|---|---|---|---|---|---|---|---|
| | | | | | | | *et al.* 2007 |
| *Chordaria linearis* | obligate | 6 | 10000000000 | clonal | aquatic | aquatic | Searles 1980, Charrier *et al.* 2007 |
| *Cladostephus verticillatus* | obligate | 8 | 125892541.2 | clonal | aquatic | aquatic | Sauvageau 1907, Charrier *et al.* 2007 |
| *Colpomenia sinuosa* | obligate | 5 | 1995262315 | clonal | aquatic | aquatic | Wynne 1972, Charrier *et al.* 2007 |
| *Conocephalum conicum* | obligate | 15 | 3162277.66 | clonal | terrestrial | aquatic | Maybrook 1914, Nishiyama 2007 |
| *Croomia pauciflora* | obligate | 42 | 25118864320 | clonal | terrestrial | aquatic | Tomlinson & Ayensu 1968 |
| *Cutleria* | obligate | 7 | 3162277660 | clonal | aquatic | aquatic | Bold & Wynne 1978, Charrier *et al.* 2007 |
| *Cyanea cyanea* | obligate | 22 | 1.00E+13 | clonal | aquatic | aquatic | Hyman 1940 |
| *Cyathodium barodae* | obligate | 13 | 50118723.36 | clonal | terrestrial | aquatic | Chavran 1937, Nishiyama 2007 |
| *Dasybranchus caducus* | obligate | 10 | 31622.7766 | clonal | NA | aquatic | Bookhaut 1957 |
| *Desmarestia antartica* | obligate | 7 | 6.31E+11 | clonal | aquatic | aquatic | Moe & Silva 1989, Charrier *et al.* 2007 |
| *Dictyosiphon hirsutus* | obligate | 6 | 39810717060 | clonal | aquatic | aquatic | Peters 1992, Charrier *et al.* 2007 |
| *Dictyostelium discoideum* | facultative | 3 | 100000 | non-clonal | terrestrial | terrestrial | Kaiser 1986, Raper 1940, Rokas 2008 |
| *Dictyostelium fasciculatum* | facultative | 2 | NA | non-clonal | terrestrial | terrestrial | |
| *Dictyostelium minutum* | facultative | 2 | 3162.27766 | non-clonal | terrestrial | terrestrial | Kaiser 198 |
| *Dictyostelium purpureum* | facultative | 2 | NA | non-clonal | terrestrial | terrestrial | Mehdiabadi *et al.* (2009 |
| *Dictyota binghamiae* | obligate | 4 | 2.51E+11 | clonal | aquatic | aquatic | Foster *et al.* 1972, Charrier *et al.* 2007 |
| *Diurodrilus westheidi* | obligate | 14 | 3548.134 | clonal | NA | aquatic | Kristensen & Niilon 1982 |
| *Ducellieria chodati* | obligate | 1 | 40 | clonal | aquatic | aquatic | Hesse *et al.* 1989 |
| *Durvillaea antartica* | obligate | 6 | 1.00E+12 | clonal | aquatic | aquatic | Naylor 1949, Charrier *et al.* 2007 |
| *Ectocarpus siliculosus* | obligate | 4 | 316227.766 | clonal | aquatic | aquatic | Knight 1931, Charrier *et al.* 2007 |
| *Elachista fucicola* | obligate | 5 | 15848931.92 | clonal | aquatic | aquatic | Koeman & Cortel-Breeman 1976, Charrier *et al.* 2007 |
| *Eudorina cylindrica* | obligate | 2 | 16 | clonal | aquatic | aquatic | Herron & Michod 2008, Hallmann 2011 |

| | | | | | | | |
|---|---|---|---|---|---|---|---|
| *Eudorina elegans* | obligate | 1 | 32 | clonal | aquatic | aquatic | Hallmann 2011 |
| *Eudorina minodii* | obligate | 1 | 32 | clonal | aquatic | aquatic | Hallmann 2011 |
| *Eudorina unicocca* | obligate | 1 | 32 | clonal | aquatic | aquatic | Yamada *et al.* 2008, Hallmann 2011 |
| *Farlowia mollis* | obligate | 7 | 3162277660 | clonal | aquatic | aquatic | Abbott 1962, Graham 1985 |
| *Fonticula alba* | facultative | 2 | | non-clonal | terrestrial | terrestrial | Brown *et al.* 2009 |
| *Fucus vesiculosis* | obligate | 7 | 3.16E+12 | clonal | aquatic | aquatic | McCully 1966, Charrier *et al.* 2007 |
| *Fuirena ciliaris* | obligate | 44 | 25118864320 | clonal | terrestrial | aquatic | Govindarajalu 1969, Graham 1985 |
| *Funaria hygrometrica* | obligate | 20 | 251188643.2 | clonal | terrestrial | aquatic | Puri 1981, Nishiyama 2007 |
| *Gloeophycus koreanum* | obligate | 12 | 63095734450 | clonal | aquatic | aquatic | Lee & Yoo 1979, Graham 1985 |
| *Gonium multicoccum* | obligate | 1 | 32 | clonal | aquatic | aquatic | Hallmann 2011 |
| *Gonium octonarium* | obligate | 1 | 32 | clonal | aquatic | aquatic | Hallmann 2011 |
| *Gonium pectorale* | obligate | 1 | 16 | clonal | aquatic | aquatic | Hallmann 2011, Herron & Michod 2008, Stein 1959 |
| *Gonium quadratum* | obligate | 1 | 16 | clonal | aquatic | aquatic | Hallmann 2011 |
| *Gonium viridistellatum* | obligate | 1 | 16 | clonal | aquatic | aquatic | Hallmann 2011 |
| *Gymnoascus reessii* | obligate | 5 | 15848.93192 | clonal | terrestrial | terrestrial | Gaetano 1986, Graham 1985 |
| *Halymenia asymmetrica* | obligate | 13 | 63095734450 | clonal | aquatic | aquatic | Gaetano 1986, Graham 1985 |
| *Haplospora globosa* | obligate | 4 | 25118864320 | clonal | aquatic | aquatic | Kuhlenkamp & Muller 1985, Charrier *et al.* 2007 |
| *Helminthostachys zeylandica* | obligate | 5 | 70794578.44 | clonal | terrestrial | aquatic | Lang 1902, Graham 1985 |
| *Heteroralfsia saxicola* | obligate | 9 | 794328234.7 | clonal | aquatic | aquatic | Kawai 1989, Charrier *et al.* 2007 |
| *Himantothallus grandifolius* | obligate | 14 | 1.58E+12 | clonal | aquatic | aquatic | Wiencke & Clayton 1990, Charrier *et al.* 2007 |
| *Hirudo medicinalis* | obligate | 26 | 19952623150 | clonal | NA | aquatic | Mann 1962 |
| *Homo sapiens* | obligate | 200 | 1.00E+14 | clonal | terrestrial | aquatic | Valentine *et al.* 1994 |
| *Hummia onusta* | obligate | 5 | 100000000 | clonal | aquatic | aquatic | Fiore 1977, Charrier *et al.* 2007 |
| *Hydra attenuata* | obligate | 15 | 63095.73445 | clonal | aquatic | aquatic | Campbell & Bode 1983, Glauber *et al.* 2010 |

| | | | | | | | |
|---|---|---|---|---|---|---|---|
| *Hymenophyllum tunbridgensis* | obligate | 15 | 7079457844 | clonal | terrestrial | aquatic | Boodle 1900, Graham 1985 |
| *Isthmoploea sphaerophora* | obligate | 3 | 15848.932 | clonal | aquatic | aquatic | Rueness 1974, Charrier *et al.* 2007 |
| *Kurogiella saxatilis* | obligate | 7 | 25118864320 | clonal | aquatic | aquatic | Kawai 1993, Charrier *et al.* 2007 |
| *Laminaria dentigera* | obligate | 14 | 1.26E+11 | clonal | aquatic | aquatic | Kain 1979, Charrier *et al.* 2007 |
| *Leathesia difformis* | obligate | 6 | 39810717060 | clonal | aquatic | aquatic | Bold & Wynne 1978, Charrier *et al.* 2007 |
| *Lemna minor* | obligate | 18 | 794328.2347 | clonal | terrestrial | aquatic | Daubs 1965, Graham 1985 |
| *Lomandra hermaphroditicum* | obligate | 36 | 35481338920 | clonal | terrestrial | aquatic | Fahn 1954, Graham 1985 |
| *Lumbricus terrestris* | obligate | 57 | 10000000000 | clonal | terrestrial | aquatic | Stephenson 1930 |
| *Mamillaria elongata* | obligate | 27 | 63095734450 | clonal | terrestrial | aquatic | Darbishire 1904 |
| *Membranoptera subtropica* | obligate | 12 | 6309573.445 | clonal | aquatic | aquatic | Schneider & Eisemann 1979, Graham 1985 |
| *Monoclea forsteri* | obligate | 13 | 3162277.66 | clonal | terrestrial | aquatic | Shuster 1984, Nishiyama 2007 |
| *Morone saxatilis* | obligate | 122 | 2.51E+11 | clonal | aquatic | aquatic | Groman 1982 |
| *Mus musculus* | obligate | 102 | 2.00E+11 | clonal | terrestrial | aquatic | Gude *et al.* 1982 |
| *Nais variabilis* | obligate | 13 | 251188.6432 | clonal | NA | aquatic | Stephenson 1908 |
| *Neodilsea natashae* | obligate | 12 | 19952623150 | clonal | aquatic | aquatic | Linstrom 1984, Graham 1985 |
| *Ophioglossum palmatum* | obligate | 14 | 6309573445 | clonal | terrestrial | aquatic | Chrysler 1941, Graham 1985 |
| *Pandorina colemaniae* | obligate | 1 | 16 | clonal | aquatic | aquatic | Bold & Wynne 1985, Hallmann 2011 |
| *Pandorina morum* | obligate | 1 | 16 | clonal | aquatic | aquatic | Bold & Wynne 1985, Hallmann 2011 |
| *Periplaneta americana* | obligate | 50 | 3162277660 | clonal | terrestrial | aquatic | Smith 1968 |
| *Petermannia cirrhosa* | obligate | 39 | 25118864320 | clonal | terrestrial | aquatic | Tomlinson & Ayensu 1968, Graham 1985 |
| *Physarum polycephalum* | facultative | 2 | 300 | clonal | terrestrial | terrestrial | Stephenson & Stempen 1994, Baldauf & Doolittle 1996, Everhart *et al.* 2008 |
| *Pinus monophylla* | obligate | 30 | 10000000000 | clonal | terrestrial | aquatic | Foster & Gifford 1974 |

| | | | | | | | |
|---|---|---|---|---|---|---|---|
| *Pisione remota* | obligate | 11 | 44668.35922 | clonal | NA | aquatic | Akesson 1961 |
| *Platydorina caudata* | obligate | 1 | 32 | clonal | aquatic | aquatic | Hallmann 2011 |
| *Pleodorina californica* | obligate | 2 | 128 | clonal | aquatic | aquatic | Hallmann 2011 |
| *Pleodorina illinoisensis* | obligate | 2 | 32 | clonal | aquatic | aquatic | Bonner 2004, Hallmann 2011 |
| *Pleodorina indica* | obligate | 2 | 128 | clonal | aquatic | aquatic | Bonner 2004, Hallmann 2011 |
| *Pleodorina japonica* | obligate | 2 | 128 | clonal | aquatic | aquatic | Hallmann 2011 |
| *Pleurobrachia* | obligate | 13 | 10000 | clonal | aquatic | aquatic | Hyman 1940 |
| *Pocheina flagellata* | facultative | 2 | NA | non-clonal | terrestrial | terrestrial | Bell & Mooers 1997, Brown *et al.* 2011 |
| *Pocheina rosea* | facultative | 2 | NA | non-clonal | terrestrial | terrestrial | Bell & Mooers 1997, Brown *et al.* 2011 |
| *Pogonatum stevensii* | obligate | 21 | 707945784.4 | clonal | terrestrial | aquatic | Chopra & Sharna 1958, Nishiyama 2007 |
| *Polysphondylium pallidum* | facultative | 2 | NA | non-clonal | terrestrial | terrestrial | Stenhouse & Williams 1980 |
| *Polysphondylium violaceum* | facultative | 2 | NA | non-clonal | terrestrial | terrestrial | Bonner 1959 |
| *Polytrichum commune* | obligate | 26 | 1000000000 | clonal | terrestrial | aquatic | Puri 1981, Nishiyama 2007 |
| *Pomatoceros triqueter* | obligate | 12 | 70794.578 | clonal | NA | aquatic | Segrove 1941 |
| *Ralfsia verrucosa* | obligate | 8 | 630957344.5 | clonal | aquatic | aquatic | Loiseaux 1968, Charrier *et al.* 2007 |
| *Saccharomyces cerevisiae* | facultative | 3 | NA | clonal | NA | terrestrial | Ratcliff *et al.* 2011, Koschwanez *et al.* 2011 |
| *Sagittaria lancifolia* | obligate | 42 | 1.00E+11 | clonal | terrestrial | aquatic | Stant 1964, Graham 1985 |
| *Salmo gairdneri* | obligate | 116 | 2.51E+11 | clonal | aquatic | aquatic | Yasutake 1983 |
| *Salpingoeca rosetta* | facultative | NA | 29 | clonal | aquatic | aquatic | Fairclough *et al.* 2010, Dayel *et al.* 2011 |
| *Sarconema scinaioides* | obligate | 13 | 2511886432 | clonal | aquatic | aquatic | Papenfuss & Edelstein 1974, Graham 1985 |
| *Schimitzia hiscockiana* | obligate | 14 | 2.00E+11 | clonal | aquatic | aquatic | Maggs & Guiry 1985, Graham 1985 |
| *Schimmelmannia dawsonii* | obligate | 11 | 2.51E+11 | clonal | aquatic | aquatic | Acleto 1972, Graham 1985 |

| | | | | | | | |
|---|---|---|---|---|---|---|---|
| *Scytosiphon lomentria* | obligate | 4 | 794328234.7 | clonal | aquatic | aquatic | Clayton 1976, Charrier *et al.* 2007 |
| *Selenipedium palmifolium* | obligate | 35 | 12589254120 | clonal | terrestrial | aquatic | Rosso 1966, Graham 1985 |
| *Sorogena stoianovitchae* | facultative | 1 | 500 | non-clonal | aquatic | terrestrial | Sugimoto & Endoh 2006, Blanton & Olive 1982, Olive & Blanton 1980 |
| *Sphacelaria bipinnata* | obligate | 9 | 1258925412 | clonal | aquatic | aquatic | Clint 1927, Charrier *et al.* 2007 |
| *Sphaerobolus stellatus* | obligate | 9 | 1258925.412 | clonal | terrestrial | terrestrial | Buller 1933, Knoll 2011 |
| *Sphagnum recurvum* | obligate | 11 | 891250938.1 | clonal | terrestrial | aquatic | Puri 1981, Nishiyama 2007 |
| *Symphyogyna brogniarti* | obligate | 13 | 446683.5922 | clonal | terrestrial | aquatic | Puri 1981, Nishiyama 2007 |
| *Syringoderma phinneyi* | obligate | 6 | 199526.2315 | clonal | aquatic | aquatic | Henry & Muller 1983, Charrier *et al.* 2007 |
| *Tetrabaena socialis* | obligate | 1 | 4 | clonal | aquatic | aquatic | Stein 1959, Herron & Michod 2008, Hallmann 2011 |
| *Volvox africanus* | obligate | 2 | 8192 | clonal | aquatic | aquatic | Herron & Michod 2008, Nozaki 2011, Hallmann 2011 |
| *Volvox aureus* | obligate | 2 | 2048 | clonal | aquatic | aquatic | Hallmann 2011 |
| *Volvox carteri* | obligate | 2 | 2048 | clonal | aquatic | aquatic | Nishii & Miller 2010, Bonner 1998, Kaiser 2001, Kirk 1999, Hallmann 2011 |
| *Volvox dissipatrix* | obligate | 2 | 16384 | clonal | aquatic | aquatic | Herron & Michod 2008, Hallmann 2011, Herron 2014 |
| *Volvox gigas* | obligate | 2 | 4096 | clonal | aquatic | aquatic | Herron & Michod 2008, Hallmann 2011, Herron 2014 |
| *Volvox globator* | obligate | 2 | 16384 | clonal | aquatic | aquatic | Herron & Michod 2008, Hallmann 2011, Herron 2014 |
| *Volvox obversus* | obligate | 2 | 2048 | clonal | aquatic | aquatic | Herron & Michod 2008, Hallmann 2011, Herron 2014 |
| *Volvox rousseletii* | obligate | 2 | 32768 | clonal | aquatic | aquatic | Herron & Michod 2008, Hallmann 2011, Herron |

| | | | | | | | |
|---|---|---|---|---|---|---|---|
| | | | | | | | 2014 |
| *Volvox tertius* | obligate | 2 | 1024 | clonal | aquatic | aquatic | Herron & Michod 2008, Hallmann 2011, Herron 2014 |
| *Volvulina boldii* | obligate | 1 | 16 | clonal | aquatic | aquatic | Herron & Michod 2008, Hallmann 2011, Herron 2014 |
| *Volvulina compacta* | obligate | 1 | 16 | clonal | aquatic | aquatic | Herron & Michod 2008, Hallmann 2011, Herron 2014 |
| *Volvulina pringsheimii* | obligate | 1 | 16 | clonal | aquatic | aquatic | Herron & Michod 2008, Hallmann 2011, Herron 2014 |
| *Volvulina steinii* | obligate | 1 | 16 | clonal | aquatic | aquatic | Stein 1958, Hallmann 2011 |
| *Wolffia arrhiza* | obligate | 5 | 10000 | clonal | terrestrial | aquatic | Luandolt 1986, Graham 1985 |
| *Wolffia microscopica* | obligate | 7 | 70794.57844 | clonal | terrestrial | aquatic | Maheshwari 1954, Graham 1985 |
| *Yamadaella cenomyce* | obligate | 7 | 3981071706 | clonal | aquatic | aquatic | Abbott 1970, Graham 1985 |
| *Yamadaphycus carnosa* | obligate | 11 | 1000000000 | clonal | aquatic | aquatic | Mikami 1973, Graham 1985 |
| *Yamagishiella unicocca* | obligate | 1 | 32 | clonal | aquatic | aquatic | Yamada *et al.* 2008, Herron & Michod 2008, Hallmann 2011 |
| *Zeacarpa leiomorpha* | obligate | 8 | 7943282347 | clonal | aquatic | aquatic | Anderson *et al.* 1988, Charrier *et al.* 2007 |
| *Zoothamnium alterans* | obligate | 4 | 141.2537545 | clonal | aquatic | aquatic | Summers 1938, Faure-Fremiet 1930 |
